# Biparental age effects in the burying beetle *Nicrophorus vespilloides*

**DOI:** 10.1101/2021.09.08.459445

**Authors:** Hilary Cope, Edward Ivimey-Cook, Jacob Moorad

## Abstract

Parental age at reproduction influences offspring size and survival by affecting prenatal and postnatal conditions in a wide variety of species, including humans. However, most investigations into this manifestation of ageing focus upon maternal age effects; the effects of paternal age and interactions between maternal and paternal age are often neglected. Furthermore, even when maternal age effects are studied, pre- and postnatal effects are confounded. Using a cross-fostered experimental design, we investigated the joint effects of paternal and pre- and postnatal maternal ages on numerous offspring outcomes in a laboratory population of a species of burying beetle, *Nicrophorus vespilloides*. When we correct our tests for significance for multiple comparisons, we found no clear evidence for any parental effect senescence acting on egg size, larval weight, or larval survival. Nor did we find a statistical effect of paternal or egg producer age on the outcomes of foster mothers as measured by weight change experienced during caregiving. These findings are consistent with recent negative results reported in a similar study of *N. vespilloides* maternal age effects while also expanding these to other offspring traits and to paternal age effects. We discuss how the peculiar life history of this species may promote selection to resist the evolution of parental age effects, and how this might have influenced our ability to detect senescence.

## Introduction

Senescence is broadly defined as the progressive loss of function due to the accumulation of damage with age and is typically associated with declining fertility and survival, known respectively as reproductive and actuarial senescence (Finch et al. 1990; Monaghan et al. 2008; Jones et al. 2014). To date, a number of comprehensive studies have reported the wide taxonomic variation in patterns of reproductive and actuarial senescence across the tree of life (Promislow 1991; Gaillard et al. 1994; Jones et al. 2008; Nussey et al. 2008; Jones et al. 2014; Lemaître and Gaillard 2017). These perspectives tend to consider only the relationship between age and outcome within individuals. However, recent attention has begun to focus upon how the age of one individual affects the phenotype of another with a specific emphasis placed upon the effects of parental age upon offspring.

Most of this research focuses upon maternal ageing, or the tendency for offspring performance to change as maternal age increases. The best-known manifestations of maternal senescence, or age-related performance declines, take the form of negative associations between maternal age two offspring outcomes: lifespan and juvenile survival. The first is known as the ‘Lansing effect’ (Lansing 1947; Comfort, A 1953; Monaghan et al. 2020); while anecdotal evidence appears to suggest that this is a common pattern across species (see Table 1 from Monaghan et al. 2020), no formal review has yet assessed its prevalence. On the other hand, maternal senescence expressed as age-related declines in juvenile survival does appear to occur more frequently than not in suitably investigated animal species groups, with the notable exception of birds (Ivimey-Cook and Moorad 2020). However, for these and other traits, great variation appears across species in both the direction and magnitudes of maternal age effects. Predictive evolutionary theory does offer some explanation for variation in maternal age effects manifested on juvenile survival (Moorad and Nussey 2016), but has not yet been generalized to explain formally the evolution of the Lansing Effect or other types of maternal senescence.

**Table 1.** Multivariate models used for data analysis

| Trait | Model number | Parental age (fixed effects)* | Random effects † |  | Replicates‡ |
| --- | --- | --- | --- | --- | --- |
| Egg length | 1 | 1. Egg-producer | 1. | Block | *n* = 2410 |
| Egg width |  | 2. Father | 2. | Egg-producer | YY = 818 |
|  |  | 3. Egg-producer X Father | ID |  | YO = 752 |
|  |  |  | 3. | Egg ID | OY = 370<br>OO = 470 |
| Larval weight at dispersal | 2 | 1. Egg-producer | 1. | Block | *n* = 697 |
|  |  | 2. Father | 2. | Carer ID | YYY = 114 |
|  |  | 3. Carer |  |  | YYO = 73 |
|  |  | 4. Egg-producer X Father |  |  | YOY = 88 |
|  |  | 5. Egg-producer X Carer |  |  | YOO = 63 |
|  |  | 6. Father X Carer |  |  | OYY = 83<br>OYO = 103<br>OOY = 90<br>OOO = 83 |
| Carer weight | 3 | 1. Egg-producer | 1. | Block | $n = 73$ |
| change |  | 2. Father |  |  | YYY = 12 |
| Larval |  | 3. Carer |  |  | YYO = 7 |
| survival to |  | 4. Egg-producer X Father |  |  | YOY = 10 |
| dispersal |  | 5. Egg-producer X Carer |  |  | YOO = 7 |
|  |  | 6. Father X Carer |  |  | OYY = 8 |
|  |  |  |  |  | OYO = 10 |
|  |  |  |  |  | OOY = 9 |
|  |  |  |  |  | OOO = 10 |

\*As carcass size is known to affect larval fitness and egg production (Müller et al. 1990; Trumbo 1990), all models also included carcass weight and carcass weight<sup>2</sup> as fixed effects. The combined weight of the two carcasses was used in the model to predict egg size, and the weight of the retained carcass was used for models predicting larval and carer outcomes.
†ID effects were nested within block effects, and the effects of egg IDs were nested within the effects of egg-producer IDs to account for possible pseudo-replication.
‡Sample sizes are given as the total number in each analysis and as each individual age combination. Ages are in the order of egg-producer, father, and carer.

Paternal age effects are much less studied than maternal age effects (Lemaître and Gaillard 2017), likely owing to the absence of post-zygotic paternal investment in most taxa which may reduce the opportunity for fathers to influence offspring (Kokko and Jennions 2008). Nonetheless, paternal age effects have been shown to exist in both human (Kong et al. 2012) and animal populations (Priest et al. 2002; Preston et al. 2015; Fay et al. 2016; Vuarin et al. 2019; Vuarin et al. 2021), with the majority showing similar detrimental effects as with maternal ageing. As a result, we may also expect comparable patterns in both juvenile survival and offspring adult lifespan across taxonomic groups, however no formal reviews have yet been conducted (although see Table 1 Monaghan et al. 2020). Our theoretical understanding of the evolution of paternal senescence lags behind that of maternal senescence, as no such models have yet been proposed.

A fuller understanding of parental age effects comes from simultaneously investigating the joint effects of both paternal and maternal age (Laaksonen et al. 2002; Priest et al. 2002; Auld and Charmantier 2011; Torres et al. 2011; Ducatez et al. 2012; Auld et al. 2013; Bouwhuis et al. 2015; Brommer et al. 2015; Preston et al. 2015; Fay et al. 2016; Whelan et al. 2016; Vuarin et al. 2021). When biparental ages are varied at least semi-independently, then maternal-by-paternal age interactions can be resolved, and this can tell us how ageing in one parent can dampen or amplify the effect of age in the other (Auld and Charmantier 2011; Ducatez et al. 2012; Auld et al. 2013; Bouwhuis et al. 2015; Whelan et al. 2016). The joint effects of biparental ageing are occasionally characterized in terms of the difference between parental ages, or PADs (parental age differences) (Fieder and Huber 2007; Helle et al. 2008; Tidière et al. 2018). The effects of PADs are related to the main and interaction effects of biparental age as *β_z,PAD_* = *β_z,M_* – *β_z,F_* – *β_z,M×F_*, where *z* is offspring performance and *β* indicates the effect upon *z* caused by differences in male (*M*) or female (*F*) age. From this, we can see that the effects of PADs are inflated relative to the differences in parental and maternal age effects when there is a negative interaction between the two, as you might expect when there is some sort of compensation for the effects of age by one parent by the other. Conversely, PAD effects are suppressed by positive interactions, such as you might expect when the ageing of one parent ‘poisons’ the offspring regardless of the age of the other parent. Despite several studies finding no significant effects of biparental ageing (Auld and Charmantier 2011; Auld et al. 2013; Bouwhuis et al. 2015), several others have highlighted the advantages of studying maternal-by-paternal age interactions. For instance, male and female ages were found to interact and advance lay date, with older males buffering the detrimental effect of reduced female experience in a population of Grey jays (*Perisoreus canadensis*, Whelan et al. 2016). In addition, in another invertebrate species, *Pieris brassicae*, male and female age at laying interacted to increase and exacerbate delays in offspring development (Ducatez et al. 2012).

In this study, we investigate the joint effects of biparental ageing on the performance *Nicrophorus vespilloides*, a species of burying beetle that demonstrates biparental care. Burying beetles are useful laboratory systems for studying parental effects because the larvae are highly amenable to cross-fostering and for this reason, the experimenter is easily able to disentangle the effects of egg-producers and carers (Lock et al. 2007: 200; Ivimey-Cook and Moorad 2018). In addition, experimental removal of one parent appears to not detrimentally affect offspring performance with females able to fully compensate for male absence (Smiseth et al. 2005). Previous work involving *N. vespilloides* has found mixed evidence of parental age effects. Some studies have found that increased maternal age at reproduction had a detrimental effect on a number of larval traits (Ward et al. 2009; Cotter et al. 2011), however a more recent paper, which disentangled egg and carer contributions, found no effect of age acting on any of the measured offspring traits (see Ivimey-Cook and Moorad 2018). Here we replicate this previous study design while expanding it to survey more offspring traits (egg length and width) and to investigate pre-natal paternal age effects (reflecting age-related differences in sperm quality or carcass preparation). This design also allows us to estimate two distinct forms of maternal-by-paternal age interaction (pre-/post-natal maternal age x paternal age). In total, we were able to investigate five different sources of parental age effects (three main effects and two interaction effects).

## Materials and Methods

### Study species

*N. vespilloides* breed on carcasses of small vertebrates and display elaborate forms of parental care (Scott 1998). Upon acquiring a carcass, a mating pair prepare it for breeding by burying it, removing all fur, scales, or feathers, rolling the carrion into a ball, and treating it with antimicrobial secretions (Smiseth et al. 2006). The female then lays eggs in the nearby soil. Newly hatched larvae aggregate on the carcass, where they both self-feed and are provisioned with pre-digested carrion by their parents, although typically the female is more involved with offspring care than the male (Smiseth et al. 2005). Parents care for their offspring until the larvae disperse from the carcass around five days after hatching. The beetles used in this experiment were bred from a large, outbred stock population maintained at the University of Edinburgh. The stock population derives exclusively from a wild population sampled from Corstophine Hill in Edinburgh, UK in 2016, and the experiment was performed in 2017. When not breeding, adults were housed individually in clear plastic containers (12 x 8 x 2cm) filled with moist soil, at 20°C, with a 16:8 light: dark photoperiod, and fed twice a week with raw organic beef.

### Experimental procedures

Female (pre- and post-natal) and male age at first reproduction were classed as either “Young” or “Old” (11-18 days or 52-65 days post-eclosion to adulthood). These age classes were chosen as they have differing levels of cumulative survival (94% and 26% respectively, Moorad*, personal communication*), presumably leading to highly disparate intensities of selection for age-specific maternal effects that should favour the evolution of maternal senescence (Moorad and Nussey 2016). Sexual maturity is reached at around 10 days post-eclosion (Lock et al. 2007). Older ages were not used here due to the scarcity of beetles surviving beyond 65 days. Varying the age at reproduction for the egg-producer, the father, and the carer, resulted in a three-factor design with eight treatment groups (Table S2). We set up a total of 154 matings; 142 of these produced eggs, and 106 produced larvae. We selected 74 of these mothers to care for a brood, of these, 73 successfully raised larvae, and one carer destroyed hers during the care period.

We weighed each outbred breeding pair (these included all combinations of male and females ages (pre- and post-natal components), see Table 1) before transferring them to a breeding box (17 x 12 x 6cm) with 2cm of moist soil containing two small mice carcasses (combined weight 17.59-25.63g) (Livefood Direct Ltd, Sheffield, UK). By providing the breeding pair with two carcasses, we expected high rates of egg and larvae production. As we describe below, one of these mice was removed from each pair to inflate larval density (see below for justifications). For each carcass pair, we marked the carcass that deviated most from the average weight of the block by removing its tail. This allowed us to reduce the among-brood variation in remaining carcass size below that of the original pool of mice. In all cases, the beetles prepared both carcasses by rolling them together into a ball.

### Egg size

We measured individual egg sizes by acquiring an image of the eggs laid three days after mating (when egg laying had ceased) through the bottom of the transparent breeding box using a Canon ConoScan 9000F Mark II (Canon Inc., Tokyo, Japan). We assigned every visible egg in each image with a unique Egg ID and measured its length and perpendicular width using 600x magnification in the dedicated software *ImageJ 1.50i* (Schneider et al. 2012). After all eggs were measured once, we repeated the process to assess the repeatability of measurements. Overall, both the length and width of 1205 eggs were measured twice (Table S1), resulting in 4820 measurements. The correlations between first and second measurements of each egg were high (Spearman’s *ρ* = +0.90 for length and +0.78 for width), indicating highly repeatable measurements.

### Cross-fostering

Females were weighed immediately after scanning eggs (three days post-mating, see section above) to establish an initial weight for each female prior to providing care to offspring. Females were then transferred along with the unmarked carcass to new breeding boxes that were absent of eggs. We disposed of the marked carcasses and the males (as we did not assess post-natal paternal age effects). Each breeding box that contained eggs was checked regularly for hatched larvae. We pooled larvae from same-aged egg-producer and paternal age groups (e.g. all offspring from young egg-producers and young fathers were pooled together) to construct mixed broods of 15 larvae. These were then provided to unrelated female carers of the either the same or differing age class. A standardized brood size of 15 was chosen as this represents a high but biologically reasonable number of larvae to be maintained on an 8.06-12.95g carcass. This represents approximately twice the density (approximately 1.1 larvae per gram of mouse rather than 0.6 larvae per gram) used in a previous *N. vespilloides* study of maternal senescence (Ivimey-Cook and Moorad 2018) and 50-100% greater density used in a *N. vespilloides* study of paternal senescence (Benowitz et al. 2013). We did this to increase the environmental stress placed upon offspring and carers, as evidence suggests that more stressful conditions can exacerbate the deleterious effects of age upon survival (Lemaître et al. 2013; Tidière et al. 2016) and reproductive output (Lemaître and Gaillard 2017). For this reason, we expected that our density manipulation would enhanced our ability to detect maternal senescence.

Larval disperse and pupate once the carcass is consumed. At this point, we counted and weighed each larva individually. The weight of larvae at dispersal is one measure believed to indicate the degree of parental reproductive investment (Ward et al. 2009). Female carers were also weighed a dispersal, and these measures were compared to those taken immediately after egg laying. The weight change of the carer over the caring period is believed to be an indicator of female investment in this species (Creighton et al. 2009; Billman et al. 2014). We then transferred post-care females to individual boxes where they were checked for death three times a week. This was to allow us to correct for selective disappearance of carers statistically by including carer age-of-death as a factor in our models.

### Statistical analyses

The experimental design required differently structured statistical models for different traits (summarized in Table 1). These were we either univariate or bivariate linear mixed-models analysed in ASReml 4.1 (Gilmour et al. 2015). The degree and nature of replication depended upon the specific traits. Egg length and width observations were made twice for each egg (2410 observations). Larval weight at dispersal was measured once for each offspring (697 observations). Carer weight change and larval survival rates were measured once per brood (73 observations). All main effects and two-way interactions between the egg-producer, paternal, and carer ages were included when biologically appropriate.

Model 3 failed to converge initially and indicated a non-positive definite variance-covariance structure for block effects. In a similar manner to Ivimey-Cook and Moorad (2018), we ran univariate analyses for each female trait with and without the random effect of block to see if we could justify dropping this random effect from the full bivariate model. Likelihood ratio tests failed to find an effect of block upon the number of larvae surviving to dispersal (*P* = 0.906), but they found a significant effect of blocks on carer weight change rerunning the full model while including block effects only for carer weight change (*P* = 0.004). We used these results to drop block effects on larval survival from our bivariate model (see Table S4 for likelihood ratio test results).

Longitudinal age-related declines in reproductive performance can be masked by selective disappearance of poor-quality individuals from the population. This effect has been reported in studies of ageing in red deer (*Cervus elaphus*) (Nussey et al. 2011), Soay sheep (*Ovis aries*) (Hayward et al. 2013) and reindeer (*Rangifer tarandus*) (Weladji et al. 2008).

However, this effect can be corrected for by including age of death as a covariate (van de Pol and Verhulst 2006; Nussey et al. 2011). Following Ivimey-Cook and Moorad (2018), we included carer age at death as a factor according to the interval during which death occurred. In this study, age had two levels: (1) between “Young” and “Old” age classes and (2) post “Old” age class. The inclusion of carer age of death into statistical models allowed us to correct for the effects of selective disappearance of carers, but not egg-producers or males, as the use of mixed broods prevented identification of genetic parents.

## Results

For brevity, only the model results for fixed age effects are provided here. Full model results are given as indicated below in the Supplemental section. *z*-scores were calculating by dividing effect sizes by standard errors, and *p* values were calculated from these. Effect sizes were divided by the median difference between the “Young” and “Old” age groups (44 days) to calculate per-day effect sizes. Significance was taken at *α* = 0.05. However, we note that the models estimated 27 effects of age (main and interaction effects), and a Bonferroni correction indicated a threshold for significance of *p* = 1.85 x 10^-3^. No estimated effect of age reached that threshold.

### Parental age effects on egg length and width

Old egg-producers and fathers produced eggs that were longer and narrower in comparison to young parents, but these effects did not reach statistical significance (Table 2). Interactions between egg-producer and father ages were negative but significantly less than zero only for effects on egg length. Differently put, matching the ages of the egg-producer and father was associated with decreased egg length. While the main effects of both parental ages were non-significant for this trait, it did appear that the matching ages of both the egg-producer and father produced significantly shorter eggs.

**Table 2.**
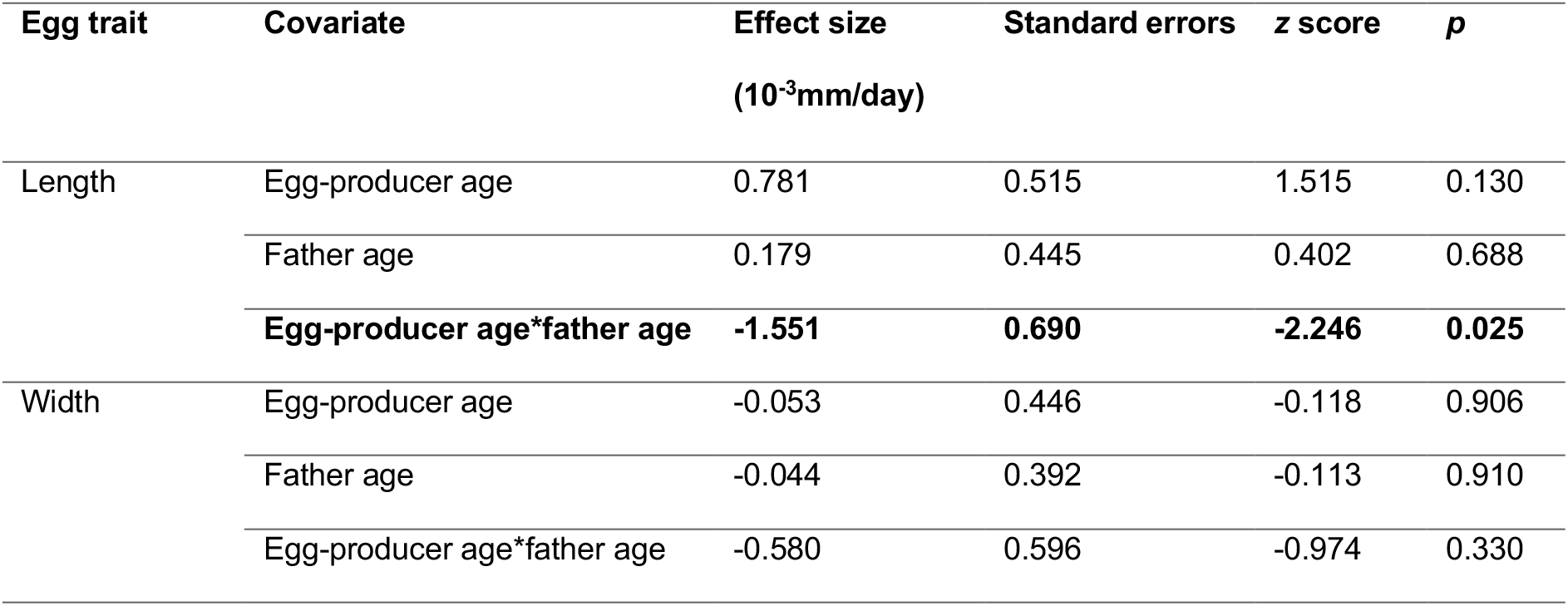
The effects of egg-producer and father age on egg size. The units of egg length and width are in mm. Effects that were significant to a threshold of *α* = 0.05 are in boldface. Full model results are given in Table S5.

| Egg trait | Covariate | Effect size<br>(10<sup>-3</sup>mm/day) | Standard errors | z score | p |
| --- | --- | --- | --- | --- | --- |
| Length | Egg-producer age | 0.781 | 0.515 | 1.515 | 0.130 |
|  | Father age | 0.179 | 0.445 | 0.402 | 0.688 |
|  | **Egg-producer age\*father age** | **-1.551** | **0.690** | **-2.246** | **0.025** |
| Width | Egg-producer age | -0.053 | 0.446 | -0.118 | 0.906 |
|  | Father age | -0.044 | 0.392 | -0.113 | 0.910 |
|  | Egg-producer age*father age | -0.580 | 0.596 | -0.974 | 0.330 |

### Parental age effects on larval weight at dispersal

Larval weight decreased with increased age of the egg producer, the father, and the carer. All two-way interaction effects were positive, larval weight increased with when both parental ages increased, but none were statistically significant (Table 3, Figure S2).

**Table 3.** The effects of egg-producer, father, and carer age on larval weight at dispersal. Full model results are given in Table S6.

| Covariate | Effect size<br>(mg/day) | Standard errors<br>(mg/day) | z score | p |
| --- | --- | --- | --- | --- |
| Egg-producer age | -0.503 | 0.316 | -1.591 | 0.112 |
| Father age | -0.216 | 0.301 | -0.718 | 0.472 |
| Carer age | -0.270 | 0.323 | -0.834 | 0.404 |
| Egg-producer age*father age | 0.235 | 0.375 | 0.627 | 0.530 |
| Egg-producer age*carer age | 0.254 | 0.375 | 0.677 | 0.498 |
| Father age*carer age | 0.075 | 0.369 | 0.204 | 0.838 |
| Age of carer at death | -0.814 | 0.800 | -1.017 | 0.309 |

### Parental age effects on carer traits, carer weight change and larval survival to dispersal

Old carers gained more weight than young carers during the parental care period. However, both old fathers and old egg-producers produced larvae that caused carers to gain less weight. All possible two-way interaction effects were positive, but none of the main or interaction effects were statistically significant to *α* = 0.05. Larval survival increased with all three main effects of age, but only the main effect of egg-producer age was statistically significant to the uncorrected level of significance. All two-way interactions were negative, with the interaction of egg-producer and carer ages significant to the uncorrected level of significance (Table 4, Figure S3-S4).

**Table 4.**
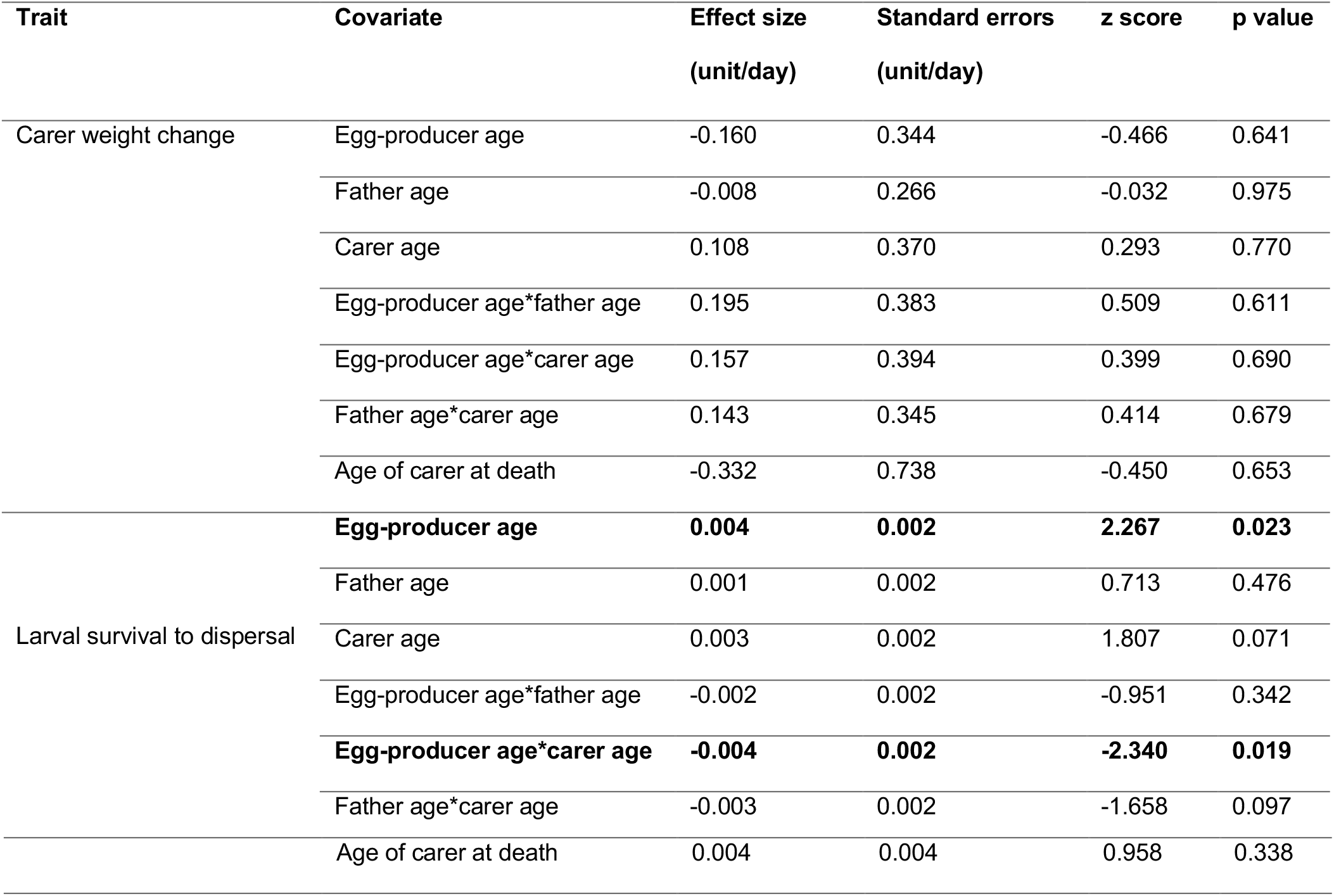
Effects of egg-producer, father, and carer age on carer weight change and larval survival. Units are *mg* for weight change measurements, and larval survival is given in terms of a fraction (#survived divided by 15 for the initial brood size). Full model results are given in Table S7.

## Discussion

There is much evidence from laboratory studies of *N. vespilloides* to suggest that maternal condition influences offspring phenotypes (Paquet and Smiseth 2017; Mattey et al. 2018; Ratz et al. 2018), but a previous study (Ivimey-Cook and Moorad 2018) found that maternal quality did not appear to change with age. Our study (which was performed in the same laboratory as all aforementioned studies) largely replicated the results reported by Ivimey-Cook and Moorad (2018), as neither detected statistical support for a general tendency towards pre- and post-natal maternal ageing to be manifested on a suite of traits related to offspring and carer fitness. Furthermore, this study suggests that this insensitivity of parental quality to changes in age also apply to pre-natal attributes of the fathers (sperm quality), to another offspring trait expected to influence fitness (egg length and width), and to a more stressful environment (high larval density). Below, we discuss how estimates from our study relate to those from other comparable studies and why the life history of this species might explain why we cannot reliably detect signatures of parental senescence.

The size of eggs reflects the degree of maternal investment made into each egg (Bagenal 1969), and because egg size appears to predict the size of newly hatched in *N. vespilloides* larvae (Monteith et al. 2012; Steiger 2013), this may be an important determinant of juvenile survival. If the resources available to a mother (or the ability of the mother to use these resources effectively) decline with age owing to senescence, then we might expect that egg size should also decline with maternal age. A previous study of hatch weight in this species found a negative effect of maternal age (Lock et al. 2007), but here we found no significant effect of either parent’s age on egg width and length. In contrast, bird studies have reported maternal age-related *increases* in egg size (Weimerskirch 1992; Bogdanova et al. 2006). However, birds and insects may operate under very different constraints in terms of maternal investment in eggs, particularly in how they are subjected to size-number trade-offs that are expected to result from limited resources. In a smaller study (700 vs 1205 eggs), Monteith *et al* (2012) reported no evidence of a negative correlation between egg size and clutch size in *N. vespilloides*, and they suggested that this may be because the costs of egg production are small in this species owing to their small size relative to the mothers. Although parental age was not directly manipulated in that study, they included maternal age as a covariate in their models and found that it had no significant relationship with egg size. Lastly, we note that increasing the size of breeding resource had a significantly negative effect on egg length (*p* = 0.020, Table S6). This could be in response to the perception by the female egg-layer that the environment was of higher quality, which would mean more resources for her larvae, and therefore less pressure to invest her own resources into her eggs. A similar pattern has been seen in several Daphnia species (Guisande and Gliwicz 1992), where egg size increases with decreasing food abundance.

Survival to dispersal is of obvious importance to fitness in this species, as successful dispersal is a necessary condition for survival to adulthood. The evolutionary theory of senescence has thus far been applied to maternal effects formally for only those traits that are important components of neonatal survival (Moorad and Nussey 2016), and this theory predicts that juvenile survival should decline at older maternal ages. However, the current study found *positive* effects of egg-producer and carer age on larval survival, although only the former was significantly different from zero. The direction and significance of both agree with results from an earlier study from the same laboratory. However, it should be noted that model specifications differed (the older study contained quadratic effects of age) owing to a different experimental design. We note that both studies detected a significant negative egg-producer-by-carer interaction effect on larval survival. With the caveat that models are structured in slightly different ways, both results suggest that larval survival increases with the age of both the egg producer and the carer, and the marginal increase in survival that a larva receives by having one old mother is suppressed if the other mother is old. It is not clear why this should be, but one possibility is that there are diminishing returns associated with benefits of ageing that are delivered to the offspring.

We found negative, but non-significant effects of all carers’, egg producers’, and fathers’ ages upon larval weight at dispersal. This trait predicts adult size, which is known to be an important factor in determining an adult’s competitive ability for securing reproductive resources (Bartlett 1988) and, therefore, for fitness. Ivimey-Cook and Moorad (2018) also reported non-significant effects of carer age, but the effects of egg-producer age were more ambiguous. In a model with first-order relationships only, the main effect of age was negative and insignificant. In a model with quadratic terms, including age-squared, the linear effect of egg-producer age was significantly greater than zero. Ward *et al*. (2009) report significantly negative effects of increased maternal age on this trait, but their experimental design conflated age with parity. Furthermore, that study did not vary the ages of carers and egg-producers independently of one-another, and the per-day effect of maternal age (−1.97 mg/day, as derived by Ivimey-Cook & Moorad (2018)) best represent the sum of pre- and post-natal age effects. If we assume no correlations between pre- and post-natal maternal effects in our study, then we can produce a similar measure by summing the two maternal effects, and we can synthesize a SE for that measure by taking the square-root of the summed squares of the individual SEs. From this, we arrive at a total maternal age effect estimate of −0.77 mg/day with a SE of 0.80 mg/day. These imply a 95%-tile interval (−2.025, 0.479) that includes the estimate from Ward *et al*. (2009). The same exercise applied to the relevant estimates taken from Ivimey-Cook and Moorad (2018) yield (−0.951, 0.777), which includes the results presented here but not the results from Ward *et al*. We note, however, that the SEs estimated here are roughly half again as large as those reported by Ivimey-Cook and Moorad (2018), and this limits the power of this exercise to compare our results for combined maternal age effects with those from other studies. However, we can infer from our results that if these combined maternal age effects are deleterious with increased age, they are likely no more extreme than those reported by Ward *et al*. (2009).

Lock *et al*. (2007) found no significant effect of the age of either the egg producer or the carer on larval weight gain. As initial larval mass is approximately two orders of magnitude smaller than weight at dispersal (Lock et al. 2007), we can take weight gain to be a reasonable proxy for the latter, and these results can be viewed as consistent with ours. The earlier study found a positive interaction effect between the ages of egg-producers and carers, which they interpreted as evidence for age-specific coadaptation for pre- and post-natal maternal effects. We also found a positive interaction, but that effect size was non-significant in our study. We note that this study used greater brood sizes than Lock *et al*. (2007) (*n* = 15 vs 9.3), which would mean more resources available to each individual offspring in the previous study.

There was no difference in the degree of weight loss between young and old carers during postnatal care. Nor did the age of the egg-producer or father appear to influence the traits of the carer. If the age of the genetic parents altered the burden placed upon the carer (by affecting the quality of the larvae, for example), then this burden was insufficient to affect carer investment enough to be detected. Further insights gained through behavioural observations could also help explain the lack of age effects on carer outcomes, particularly if parental care behaviours are changing as a result of increasing egg-producer and father age.

Given that female investment into reproduction may be influenced by age-dependent characteristics of the partner, it is important to consider any paternal age-effects on reproductive performance, but studies that investigate these are rare and show a diversity of results. We found that paternal age has no detectable effect on offspring quality in *N. vespilloides;* and this result is supported by Ward *et al*. (2009) who found no detectable effect upon mass at dispersal in this species. Fox *et al*. (2003) also found no evidence that paternal age affected any offspring traits in a study of the seed beetle *Callosobruchus maculatus*. However, Fay *et al*. (2016) found that the age of fathers in the wandering albatross, *Diomedea exulans*, significantly influenced offspring performance, with increasing paternal age leading to declines in juvenile survival. Paternal age in humans was shown to be the predominant factor behind the exponential increase in the number of *de novo* mutations occurring within children (Kong et al. 2012), and increased paternal age in the houbara bustard, *Chlamydotis undulata*, was found to decrease sperm quality and hatching success and to slow neonatal development (Preston et al. 2015). Conversely, male age in a different bird species, *Colaptes auratus*, was associated with increased clutch size, earlier laying and greater fledging success (Wiebe 2018). Results from the present study appear to support the notion that no clear patterns of directional effects exist.

The effects of differences between the age of father and mothers (*parental age differences*, or PADs) have received interest in studies of human reproduction, where they appear to predict family size (Fieder and Huber 2007; Helle et al. 2008). Tidière *et al*. (2018) recently pointed out, however, that the effects of PADs are statistically entangled with the main effects of parental age, and for this reason they recommend that single parent ages be used as covariates in analyses of PAD effects. We agree with this reasoning, but Tidière *et al*. (2018) leave out a statistically explicit definition of PADs that we provide here (*β_z,PAD_* = *β_z,M_* – *β_z,F_* – *β_z,M×F_*), where main and interaction effects can be estimated simultaneously by GLMs, as we have done here. As we have noted, we detected effects of interactions between father and egg-producer upon egg length in *N. vespilloides*. In this case, effect sizes estimates given in Table 3 define the PAD effect to be equal to 0.949 x 10^-3^mm/day. Many other studies estimate parental-age-interaction effects (Laaksonen et al. 2002; Auld et al. 2013; Brommer et al. 2015; Whelan et al. 2016), and these can be converted to define PAD effects in the same way to provide a clearer perspective of causality.

Selective disappearance has been shown to be an important contributor to population-level ageing patterns in wild vertebrate systems (Nussey et al. 2006; Hayward et al. 2013), but it is seldom investigated in laboratory studies of ageing. Our study controlled for selective disappearance of carers, but we found no evidence that mortality biased parental age effects. Ivimey-Cook and Moorad (2018) also found no evidence for selective disappearance of carers. Unfortunately, neither *N. vespilloides* study could investigate selective disappearance of egg producers or father, as both experiments used mixed broods of larvae, and identification of the genetic parents of larvae is not possible. Using intact broods of cross-fostered larvae, where larval siblings were cross-fostered to a non-related carer, would have allowed the age of death of all three parents to be accounted for. However, a pilot study suggested that this would be extremely difficult owing to both asynchronous hatching of larvae within a brood and the high frequency of broods with small numbers of larvae. A further study using a larger combined carcass size would, in theory, increase the number of eggs laid and larvae produced by each breeding pair, and allow more successful cross-fostering of intact broods. In this way, potential bias arising from selective disappearance of the other parents could be assessed, too.

A recent review of the literature that examines maternal age effects on early survival (Ivimey-Cook and Moorad 2020) found that among widely-studied taxa, insect species feature the strongest tendencies towards maternal effect senescence (17 of 26 species exhibited senescence). As our results show that maternal effect senescence is not manifested on offspring traits; this suggests that *N. vespilloides* is unusual. This species has a peculiar life history that we suggest indicates an evolutionary cause for slowed (and, thus, harder to detect) rates of maternal effect senescence. Evolutionary theory demonstrates that selection for the maternal effects upon neonatal survival is proportional to the age distribution of maternal ages (Moorad and Nussey 2016). It follows that when reproduction is highly focused upon a small range of ages, then the decline in selection for maternal effects that follows the onset of reproduction will be dramatic, and evolution will favour extreme rates of ageing. When reproduction is spread over many ages, the opposite is expected: ageing should be gradual. Reproduction in *N. vespilloides* depends upon the availability of vertebrate carcasses, and this is a rare and unpredictable resource (Scott 1998). As a result, we can expect high variation in the age of first reproduction and in the time between successive reproductive events. These features will cause the shape of the frequency distribution of maternal ages at birth to favour a late maximum (the age of strongest selection) and subsequent slow declines afterwards (a slow rate of relaxing selection). The latter feature will promote strong selection to resist senescence in maternal age effects, which is consistent with the observations made here for larval survival. Whilst the evolutionary theory makes explicit predictions regarding only maternal effects on neonatal survival, it seems reasonable to expect that similar predictions would apply both to other offspring traits and to paternal age effects, but the future models should be developed to generalize the theory to consider these other traits and factors.

