## Supplementary material for "Biparental age effects in the burying beetle *Nicrophorus vespilloides*": Table S1

**Table S1.** Eggs measured in each treatment

| Treatment (Egg-producer age, father age) | Number of eggs measured |
|---|---|
| Young, young | 409 |
| Young, old | 376 |
| Old, young | 185 |
| Old, old | 235 |
| Total | 1205 |

**Table S2.** Number of cross-fostered broods achieved, and the total number of surviving larvae at dispersal in each treatment

| Treatment (Egg-producer age, father age, carer age) | Number of broods | Number of larvae |
|---|---|---|
| Young, young, young | 12 | 114 |
| Young, young, old | 7 | 73 |
| Young, old, young | 10 | 88 |
| Young, old, old | 7 | 63 |
| Old, young, young | 8 | 83 |

| | | |
|---|---|---|
| Old, young, old | 10 | 103 |
| Old, old, young | 9 | 90 |
| Old, old, old | 10 | 83 |
| Total | 73 | 697 |

**Table S3.** Summary of Akaike information criterion comparison for model selection

| Traits | Null | First-order | First- and second-order |
|---|---|---|---|
| Egg length | -4077.287 | -4074.113 | -4076.405 |
| Egg width | -5804.061 | -5802.894 | -5802.615 |
| Larval weight at dispersal | -3025.628 | -3021.442 | -3016.943 |
| Carer weight change | -293.690 | -289.815 | -284.576 |
| Larval survival to dispersal | 360.484 | 363.968 | 358.372 |

**Table S4**. Likelihood ratio tests for bivariate carer-level models with and without block effects

| Trait | Log-likelihood with block | Log-likelihood without block | *D* | p value |
|---|---|---|---|---|
| Carer weight change | 188.293 | 184.108 | 8.370 | **0.00381** |
| Larval survival to dispersal | 108.090 | 108.097 | 0.014 | 0.906 |

**Table S5.** Parameter estimates from the full bivariate model assessed at the level of the egg.

| Trait | Covariate | Effect size ($10^{-3}$ mm/day) | Standard errors | z score | p -value |
|---|---|---|---|---|---|
| Egg length | **Intercept** | **56.522** | **0.306** | **184.633** | **<0.001** |
| | Egg-producer age | 0.781 | 0.515 | 1.515 | 0.130 |
| | Father age | 0.179 | 0.445 | 0.402 | 0.688 |
| | **Egg-producer age*father age** | **-1.551** | **0.69** | **-2.246** | **0.0247** |
| | **Carcass weight** | **-0.771** | **0.333** | **-2.320** | **0.0203** |
| | Carcass weight$^2$ | -0.622 | 0.645 | -0.964 | 0.335 |
| Egg width | **Intercept** | **29.614** | **0.271** | **109.404** | **<0.001** |
| | Egg-producer age | -0.0528 | 0.446 | -0.118 | 0.906 |
| | Father age | -0.0442 | 0.392 | -0.113 | 0.910 |
| | Egg-producer age*father age | -0.580 | 0.596 | -0.974 | 0.330 |
| | Carcass weight | 0.0441 | 0.284 | 0.155 | 0.877 |
| | Carcass weight$^2$ | -0.456 | 0.553 | -0.825 | 0.409 |

**Table S6.** Parameter estimates from the full univariate linear model assessed at the level of the larvae

| Trait | Covariate | Effect size (mg/day) | Standard errors | z score | p |
|---|---|---|---|---|---|
| Larval weight at dispersal | **Intercept** | **5.127** | **0.220** | **23.279** | **<0.001** |
| | Egg-producer age | -0.503 | 0.316 | -1.591 | 0.112 |
| | Father age | -0.216 | 0.301 | -0.718 | 0.472 |
| | Carer age | -0.270 | 0.323 | -0.834 | 0.404 |
| | Egg-producer age*father age | 0.235 | 0.375 | 0.627 | 0.530 |
| | Egg-producer age*carer age | 0.254 | 0.375 | 0.677 | 0.498 |
| | Father age*carer age | 0.0752 | 0.369 | 0.204 | 0.838 |
| | Age of carer at death | -0.814 | 0.800 | -1.017 | 0.309 |
| | **Carcass weight** | **0.585** | **0.236** | **2.476** | **0.0133** |
| | Carcass weight$^2$ | 0.383 | 0.347 | 1.103 | 0.270 |

**Table S7.** Parameter estimates from the full bivariate model assessed at the level of the carer.

| Trait | Covariate | Effect size (unit/day) | Standard errors | z score | p |
|---|---|---|---|---|---|
| Carer weight change | Intercept | -0.530 | 0.325 | -1.633 | 0.102 |
| | Egg-producer age | -0.160 | 0.344 | -0.466 | 0.641 |
| | Father age | -0.00843 | 0.266 | -0.0316 | 0.975 |

| | | | | | |
|---|---|---|---|---|---|
| | Carer age | 0.108 | 0.370 | 0.293 | 0.770 |
| | Egg-producer age*father age | 0.195 | 0.383 | 0.509 | 0.611 |
| | Egg-producer age*carer age | 0.157 | 0.394 | 0.399 | 0.690 |
| | Father age*carer age | 0.143 | 0.345 | 0.414 | 0.679 |
| | Age of carer at death | -0.332 | 0.738 | -0.450 | 0.653 |
| | Carcass weight | -0.198 | 0.226 | -0.874 | 0.382 |
| | Carcass weight$^2$ | 0.475 | 0.326 | 1.459 | 0.145 |
| **Larval survival to dispersal** | **Intercept** | **0.0130** | **0.00110** | **11.797** | **<0.001** |
| | **Egg-producer age** | **0.00363** | **0.00160** | **2.267** | **0.0234** |
| | Father age | 0.00110 | 0.00155 | 0.713 | 0.476 |
| | Carer age | 0.00296 | 0.00164 | 1.807 | 0.0708 |
| | Egg-producer age*father age | -0.00181 | 0.00190 | -0.951 | 0.342 |
| | **Egg-producer age*carer age** | **-0.00447** | **0.00191** | **-2.340** | **0.0193** |
| | Father age*carer age | -0.00313 | 0.00189 | -1.658 | 0.0973 |
| | Age of carer at death | 0.00397 | 0.00414 | 0.958 | 0.338 |
| | **Carcass weight** | **-0.00287** | **0.00120** | **-2.401** | **0.0164** |
| | Carcass weight$^2$ | -0.00197 | 0.00176 | -1.119 | 0.263 |

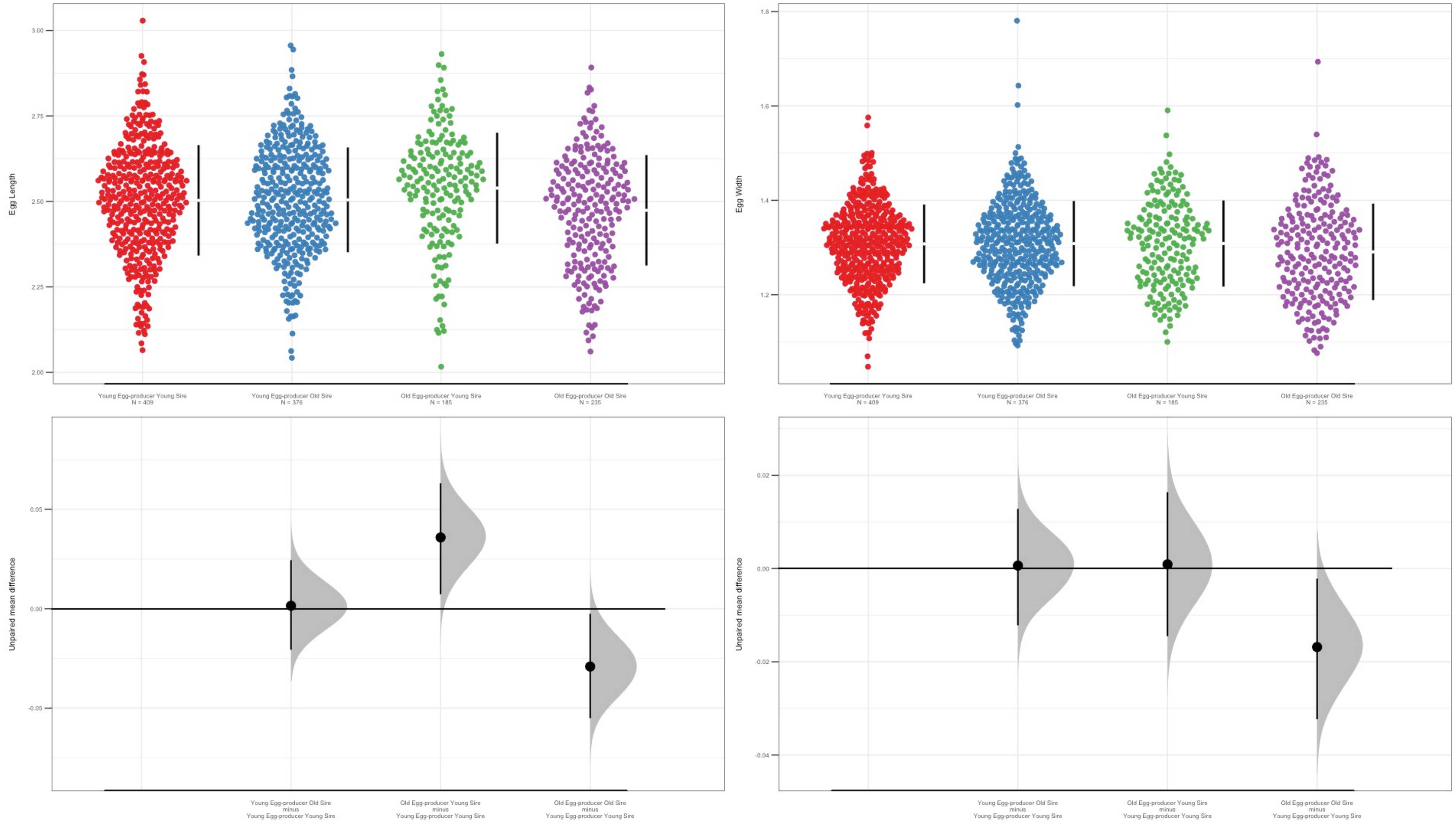


**Figure S1.** Effects of egg-producer and male age on egg length (left) and width (right). Each colour represents a different combination of egg-producer and male age. Each panel shows the distribution of egg measurements with mean and bootstrapped 95% confidence intervals in the top plot. The bottom plot shows the mean difference between the reference level (Young egg-producer and young sire) and other combinations of male and egg-producer age.

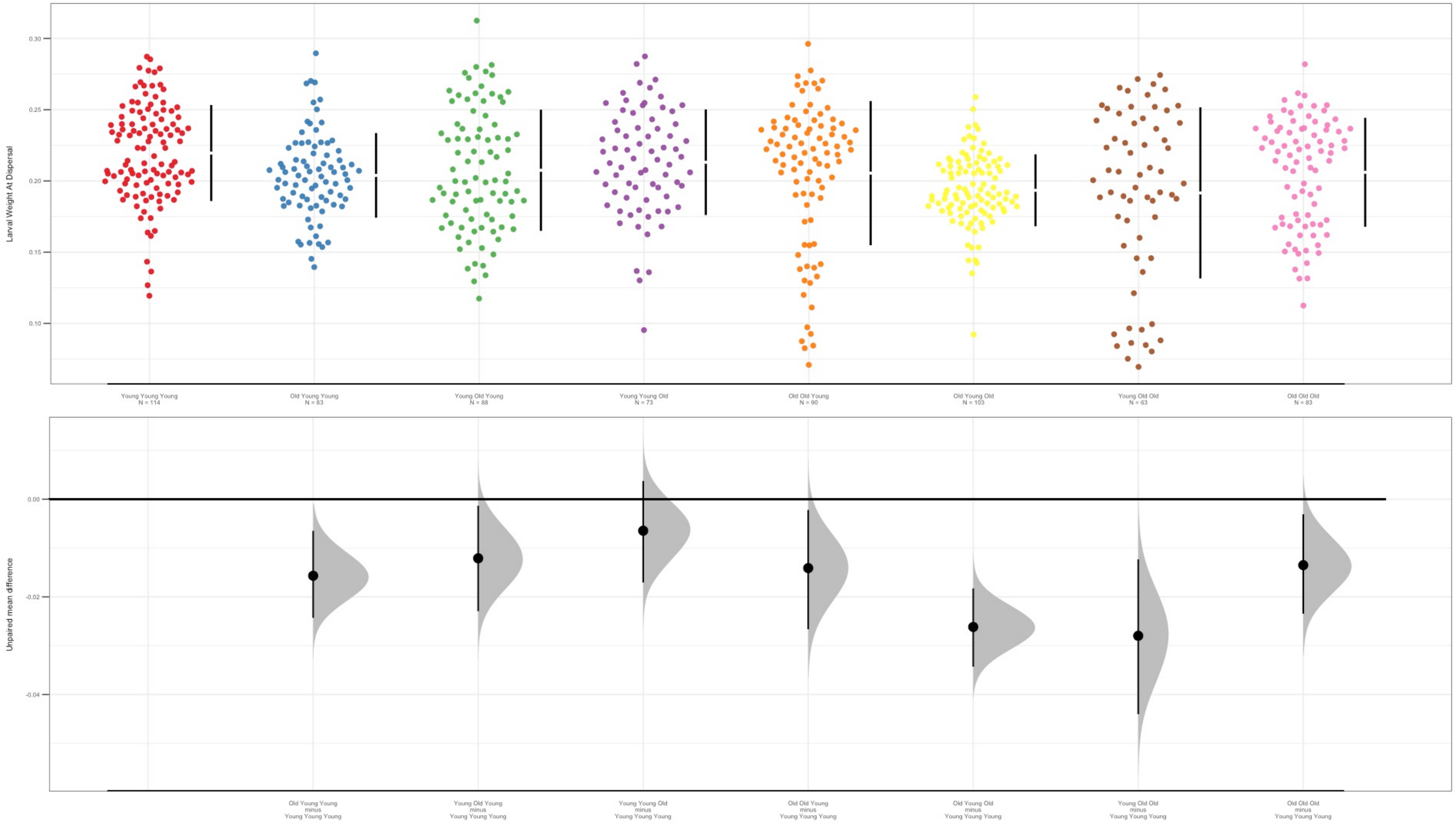


**Figure S2.** Effects of egg-producer, male and carer age on larval weight at dispersal. Each colour represents a different combination of egg-producer, male and carer age. Each panel shows the distribution of larval dispersal weights with mean and bootstrapped 95% confidence intervals in the top plot. The bottom plot shows the mean difference between the reference level (Young egg-producer, young sire and young carer) and other combinations of egg-producer, male and carer age.

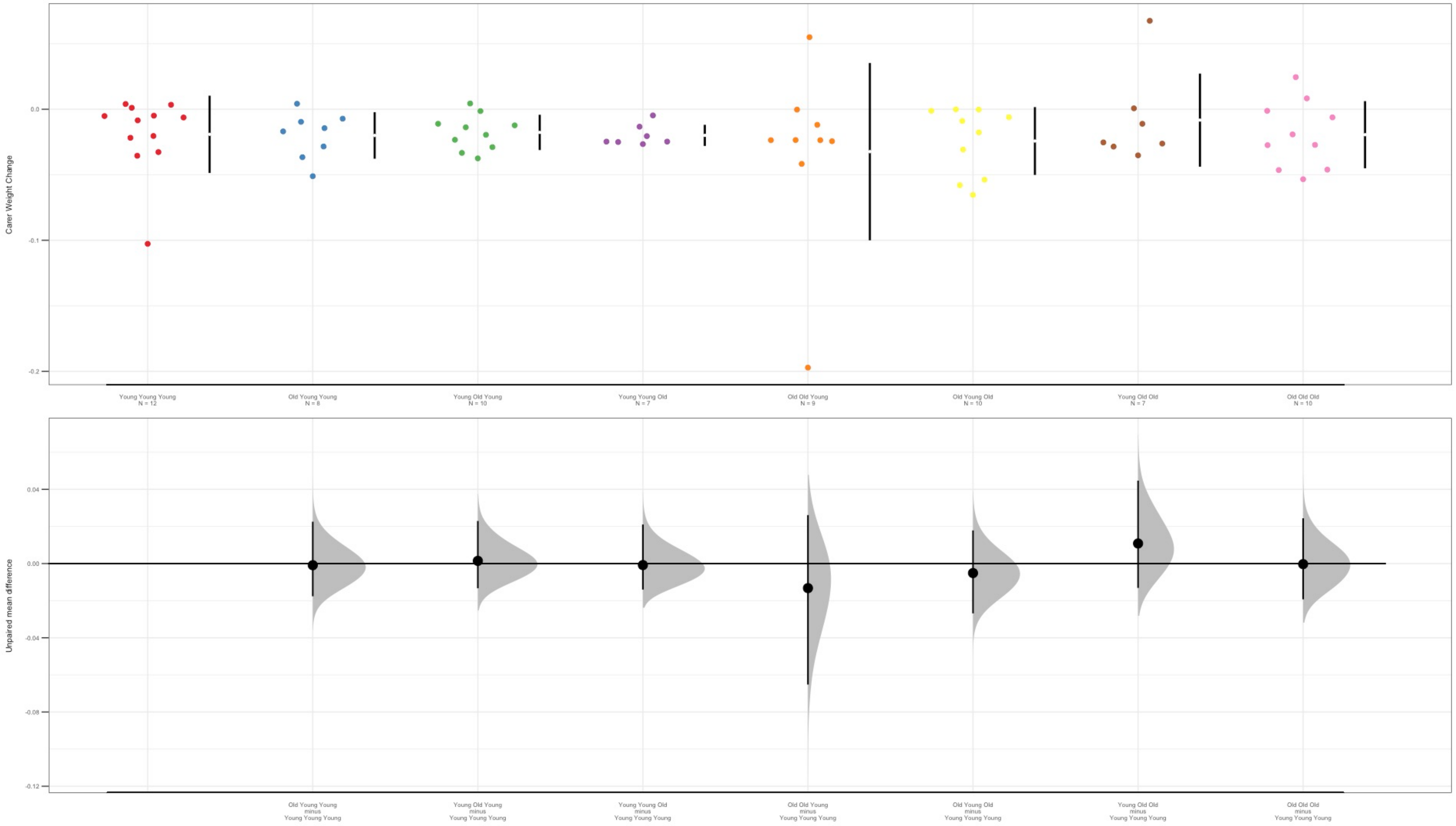


**Figure S3.** Effects of egg-producer, male and carer age on number of larvae surviving to dispersal. Each colour represents a different combination of egg-producer, male and carer age. Each panel shows the distribution of surviving larval number for each female with mean and bootstrapped 95% confidence intervals in the top plot. The bottom plot shows the mean difference between the reference level (Young egg-producer, young sire and young carer) and other combinations of egg-producer, male and carer age.

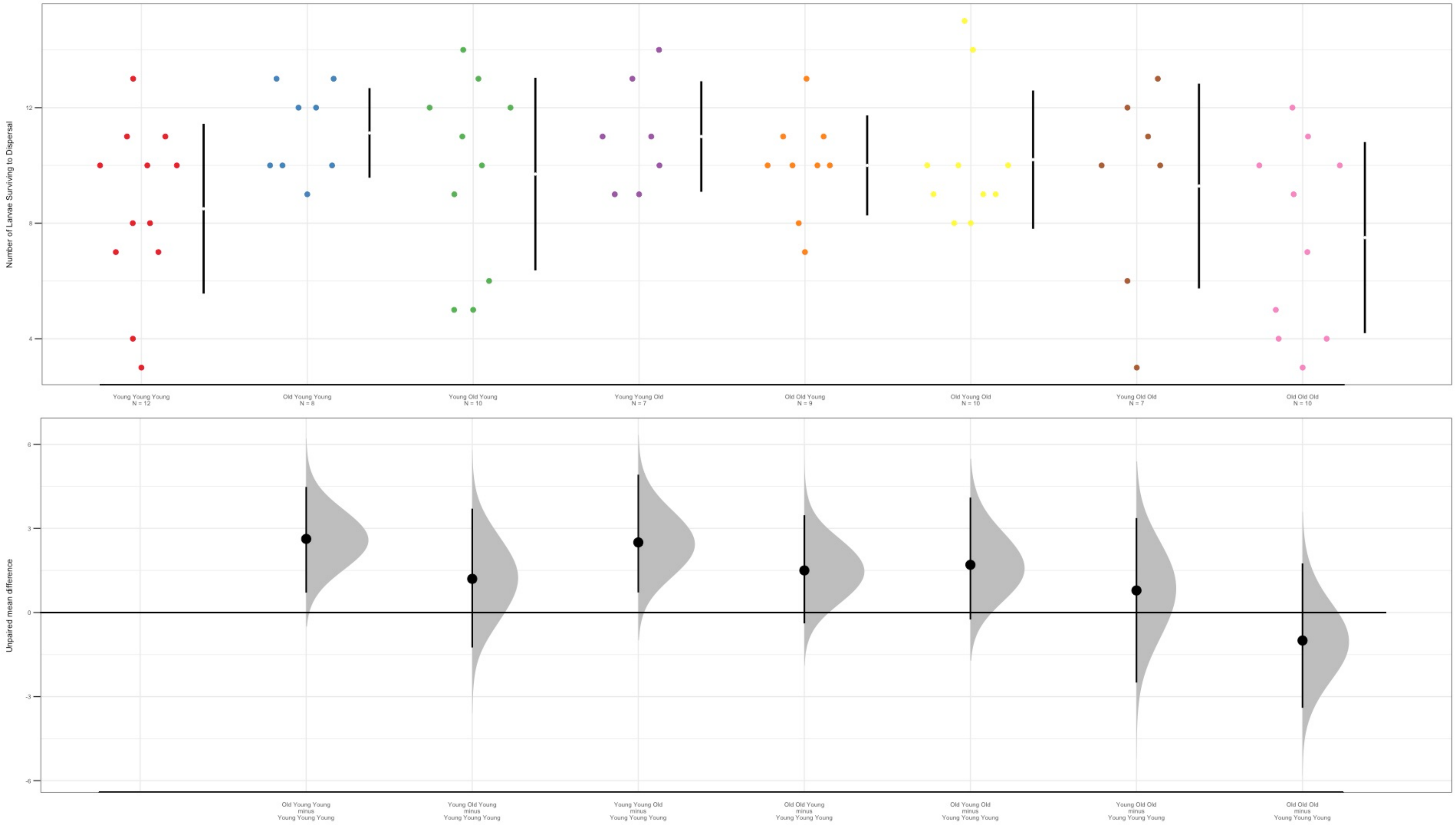


**Figure S4.** Effects of egg-producer, male and carer age on the number of larvae surviving to dispersal. The boxplots show the minimum and maximum values, along with the median and interquartile range for all treatment groups.
